# Characterisation and sequence mapping of large RNA and mRNA therapeutics using mass spectrometry

**DOI:** 10.1101/2022.02.14.480356

**Authors:** C.J. Vanhinsbergh, A. Criscuola, J. Sutton, K. Murphy, A.J.K Williamson, K. Cook, M.J. Dickman

## Abstract

Large RNA including messenger RNA (mRNA) has emerged as an important new class of therapeutic. Recently this has been demonstrated by two highly efficacious vaccines based on mRNA sequences encoding for a modified version of the SARS-CoV-2 spike protein. There is currently significant demand for the development of new and improved analytical methods for the characterization of large RNA including mRNA therapeutics. In this study we have developed an automated, high throughput workflow for the rapid characterisation and direct sequence mapping of large RNA and mRNA therapeutics. Partial RNase digestions using RNase T1 immobilised on magnetic particles was performed in conjunction with high resolution liquid chromatography mass spectrometry analysis. Sequence mapping was performed using automated oligoribonucleotide annotation and identifications based on MS/MS spectra. Using this approach >80% sequence of coverage of a range of large RNAs and mRNA therapeutics including the SARS Co-V2 spike protein was obtained in a single analysis. The analytical workflow, including automated sample preparation can be completed within 90 minutes. The ability to rapidly identify, characterise and sequence map large mRNA therapeutics with high sequence coverage provides important information for identity testing, sequence validation and impurity analysis.

## 1. INTRODUCTION

Large RNA including mRNA has recently emerged as a new class of therapeutic as demonstrated by the development and approval of two highly efficacious vaccines based on mRNA sequences encoding for a modified version of the SARS-CoV-2 spike protein ^1-2^. During the enzymatic manufacturing process of mRNA therapeutics, incomplete mRNA products are generated in conjunction with other potential impurities such as dsRNA. Furthermore, during manufacturing and storage, RNA and RNA therapeutics can be degraded by exposure to heat, hydrolysis, oxidation, light and ribonucleases. The development of analytical methods for the analysis of mRNA therapeutics is critical to underpin manufacturing development. Analytical methods are also required to assess batch to batch manufacture, process repeatability as well as the quality of mRNA produced. Furthermore, validated analytical methods are required to support the relevant phase of clinical development, regulatory submission requirements or to support ongoing quality control of the approved product ^3^. Current analytical methods available to characterise RNA therapeutics are limited and the development of methods for the analysis of large RNA (>1000 nucleotides) including mRNA vaccines is challenging. Therefore, there is currently significant demand for the development of new and improved analytical methods for the characterization of mRNA therapeutics.

Mass spectrometry based methods offer a powerful approach to analyse RNA. RNase mapping methods have been developed previously and used in a wide variety of applications including RNA sequence mapping and identification of RNA post transcriptional modifications ^4-13^. Enzymatic digestion of the RNA using ribonucleases such as RNase T1 or A generates smaller oligoribonucleotides that are more amenable for chromatographic separation and intact mass measurements ^14-18^ additional sequence information of the oligoribonucleotides can be obtained using tandem mass spectrometry (MS/MS) ^19-20^. However, the use of high frequency RNase enzymes for RNA sequence mapping of long RNA and mRNA therapeutics results in the production of a large number of small oligoribonucleotides which map to multiple different locations throughout the RNA sequence and therefore do not generate unique sequences for sequence mapping. Furthermore, the analysis of RNase sequence mapping MS data is challenging and currently there are limited dedicated software tools available which often require significant manual annotation.

Recent approaches have further developed the use of RNase mass mapping for the characterization of mRNA and sgRNA ^21-22^. RNase digestion of the mRNA was performed using RNases such as RNase T1 in conjunction with alternative RNase enzymes, including MazF, for RNase mass mapping approaches ^21^. Utilising multiple orthogonal enzymes increased the sequence coverage of the long mRNA compared to using single RNase enzymes and demonstrates the application of LC-MS/MS as an important approach for the analysis of the of mRNA therapeutics. Similar parallel nuclease digestions using alternative RNases have also recently been used to sequence map sgRNA ^22^.

In this study we have developed a rapid, automated method for the direct characterisation and comprehensive sequence mapping of large RNA and mRNA therapeutics. Partial RNase digestions using RNase T1 immobilised on magnetic particles were performed prior to high resolution LC MS/MS. The novel use of controlled partial digestion was desigened to induce missed cleavages in order to create longer fragments that could be uniquely matched to the original RNA sequence. Oligoribonucleotide sequence identification was performed using newly developed data analysis software that enables the automatic identification of multiple missed cleavages in conjunction with accurate intact mass analysis and MS/MS fragmention spectra. These were then used to generate a high coverage sequence map based on the corresponding RNA sequence. Using this novel workflow >80% sequence coverage of a range of large RNAs and mRNA therpeutics, inlcuding mRNA for the SARS CoV-2 spike protein, was generated from a single partial RNase T1 digest.

## 2. EXPERIMENTAL SECTION

### 2.1. Chemicals

Water (UHPLC MS grade, Thermo Scientific), Acetonitrile (UHPLC MS grade, Thermo Scientific), 1,1,1,3,3,3,-Hexafluoro-2-propanol (HFIP, >99.8% Fluka LC MS grade), Triethylammonium acetate (TEAA, Sigma), Triethylamine (TEA, 99.7% extrapure Fisher Scientific). SMART Digest Bulk magnetic RNase T1 Kit, (Thermo Scientific).

### 2.2. *In vitro* transcription (IVT) of RNA

mRNA synthesis via *in vitro* transcription was performed using linearized plasmid DNA using High Scribe T7 Polymerase HiScribe™ T7 High Yield RNA Synthesis Kit (New England Biolabs): 100 mM NTPs, 1x reaction buffer, 1 µg DNA template and 2 µl HiScribe™ T7 polymerase in 20 µl RNase-free water. eGFP mRNA was prepared using a DNA template containing the open reading frame flanked by the 5′ and 3′ untranslated regions (UTR) and a poly-A tail. SARS CoV-2 Spike protein mRNA was prepared using a DNA template containing the open reading frame flanked by the 5′ and 3′ UTR. Following IVT, template DNA was removed by the addition of DNase I and RNA was purified using silica columns as previously described ^23^. RNA concentrations were determined using a NanoDrop™ 2000c spectrophotometer (ThermoFisher Scientific) by absorbance at 260 nm normalized to a 1.0 cm (10.0 mm) path. Additional analysis of the RNA was subsequently performed using ion-pair reverse phase chromatography to assess the purity of the RNA. CleanCap Fluc mRNA and CleanCap 5 Fluc mRNA (5-methoxyuridine) were purchased from TriLink Biotechnologies.

### 2.3. Ion Pair-Reverse Phase High Performance Liquid Chromatography (IP-RP HPLC) of intact mRNA

Samples were analysed by IP-RP-HPLC on a U3000 HPLC system using a DNAPac RP (150 mm x 2.1 mm I.D. Thermo Fisher Scientific). Chromatograms were generated using UV detection at a wavelength of 260 nm. The chromatographic analysis was performed using the following conditions: Buffer A 100 mM triethylammonium acetate (TEAA) pH 7.0; Buffer B 0.1 M TEAA, pH 7.0 containing 25% acetonitrile. RNA was analysed using the following gradient. Gradient starting at 22% buffer B to 27% in 2 minutes, followed by a linear extension to 62% buffer B over 15 minutes, then extended to 73% buffer B over 2.5 minutes at a flow rate of 0.4 ml/min at 50 °C with UV detection at 260 nm.

### 2.4. RNase sequence mapping

Partial RNase digestions were performed using 20-40 µg of RNA incubated with 1.25-5 µl of immobilised RNase T1 at either 60 °C or 37 °C for 5-15 min in a volume of 50 µl of the SMART digest RNase buffer. Reactions were stopped by the magnetic removal of the immobilised RNase T1. Automated RNase digestions were performed on a KingFisher Duo Prime system (Thermo Scientific) using BindIt™ software (version 4.0) to control the KingFisher Duo Prime system. A 96-deepwell plate was set up with 50 μl of SMART digest RNase buffer containing 20-40 µg of RNA samples in Row A and 2.5-5 µl RNase T1 immobilized on magnetic beads within 50 μl of SMART digest RNase T1 buffer in Row G. The KingFisher was programmed to transfer RNase T1 immobilized magnetic particles to Row A, to digest the RNA at 37 °C for 5-15 minutes. Sedimentation of beads was prevented by repeated insertion of the magnetic comb using the mixing speed setting “Fast”. Immediately after incubation, the magnetic beads were collected and removed from the reaction and the digest solution was actively cooled to 15 °C. Complete RNase digests were performed using 10-20 µg of RNA with the addition of 100 U RNase T1 (Thermo Fisher Scientific) at 37 °C for 4 hrs in 0.1M TEAA. Subsequently, 10-20 µg of digested RNA was analysed using LC MS/MS.

RNA digests were analysed by IP-RP-HPLC on a Vanquish binary gradient UHPLC system (Thermo Fisher Scientific) using a DNAPac RP (300 mm x 2.1 mm I.D. Thermo Scientific). LC buffer A: 0.2% Triethylamine (TEA) 50 mM 1,1,1,3,3,3,-Hexafluoro-2-propanol. LC buffer B: 0.2% Triethylamine (TEA) 50 mM 1,1,1,3,3,3,-Hexafluoro-2-propanol, 20% ^v^/_v_ acetonitrile. Starting with 2% buffer B, a linear extension to 25% B in 40 mins, at 60°C, at a flow rate of 200 µl min^-1^, using UV detection at a wavelength of 260 nm. Mass Spectrometry analysis was performed using an Orbitrap Exploris 240 LC MS instrument (Thermo Fisher Scientific). Data acquisition was performed using DDA in full scan negative mode, scanning 450 to 3000 m/z, with an MS1 resolution of 120,000 and normalized Automatic Gain Control (AGC) target of 200%. MS1 ions were selected for higher energy collisional dissociation (HCD). MS2 resolution was set at 30,000 with the AGC target of 50%, with isolation window of 4 m/z and scan range of 150-2000 m/z, normalized stepped collision energy 15, 18, 21.

### 2.5. LC MS/MS Data Analysis

Data analysis was performed in BioPharma Finder v5.0 (Thermo Fisher Scientific). Data analysis used the basic default method in the oligonucleotide sequencing module. To identify large fragment ions, the maximum oligonucleotide mass was set to 25,000 Da, minimum confidence at 0.5 and mass accuracy at 10 ppm. The ribonuclease selection was set to RNase T1, specificity level set at ‘strict’ and phosphate location was set at ‘none’. Phosphorylation and cyclic phosphorylation were set as variable modifications of the 3’ terminal in the sequence manager containing the RNA sequence. Random RNA sequences of the same length and GC content were included in the sequence manager in addition to the correct RNA sequence. For data processing and review, additional filters were included to discount nonspecific identifications, to contain MS/MS data in each identification with a confidence score above 90%, best overall structural resolution below 2.0 and delta mass accuracy 20 ppm. All oligonucleotide identifications from BioPharma Finder are shown in Supplementary Table T1.

## 3. RESULTS AND DISCUSSION

### 3.1. Development of a workflow for the sequence mapping of mRNA and long RNA using LC MS/MS

Complete digestion of large RNA including mRNA therapeutics using RNases such as RNase T1/A results in the production of a large number of small oligoribonucleotides which map to multiple different locations throughout the RNA sequence and therefore do not generate unique sequences for sequence mapping ^14^. Furthermore, many oligoribonucleotide fragments generated are isobaric and cannot be identified based on high resolution accurate mass analysis (HRAM) alone. Therefore, RNase sequence mapping in conjunction with complete digestion of large RNA using RNases such as RNase T1/A results in limited sequence coverage. To overcome these limitations, we developed a simple workflow using partial RNase T1 digests in conjunction with high resolution LC MS/MS and automated software tools for RNA sequence mapping. Partial RNase digests have previously been used in conjunction with complete RNase digestions to determine the sequence of tRNA^Val^ from *Torulopsis utilis* in conjunction with HPLC and complete digestion of the fragments using snake venom phoshodiesterase to the component nucleotides ^24^.

To achieve reproducible partial RNase digestions is challenging, in this study reactions were performed using specifically developed RNase T1 immobilised on magnetic particles. This allowed simple control of the enzymatic reaction, which could be effectively stopped by simply removing the magnetic particles after a short, defined period of time. Utilising RNase T1 immobilised on magnetic beads also enables automation of the workflow which was performed both manually and further developed on an automated robotic liquid handling system which enabled automation of the RNase T1 digest and sample preparation for direct analysis using LC MS/MS. Furthermore, the use of RNase T1 immobilised on magnetic particles also prevents build-up of RNase T1 on the HPLC column, from standard in-solution digests, which can potentially further digest the RNA during chromatographic separation ^18^ which in this case would potentially further digest the oligoribonucleotides generated from the partial RNase digest. This is important to avoid, as the partial digestion is designed to release larger oligoribonucleotide fragments, which when sequenced will locate to a unique position within the mRNA. Following partial RNase T1 digestion, the oligoribonucleotides were separated using ion pair reverse phase HPLC (IP RP HPLC) in conjunction with mass spectrometry analysis. Oligoribonucleotide identifications were performed using newly developed automated data analysis software which is able to identify oligoribonucleotides based on their accurate mass in conjunction with the MS/MS fragmentation spectra and map the corresponding oligoribonucleotide sequences to the known RNA sequence. A schematic illustration of the total mRNA sequencing workflow is shown in Figure 1.

**Figure 1.**
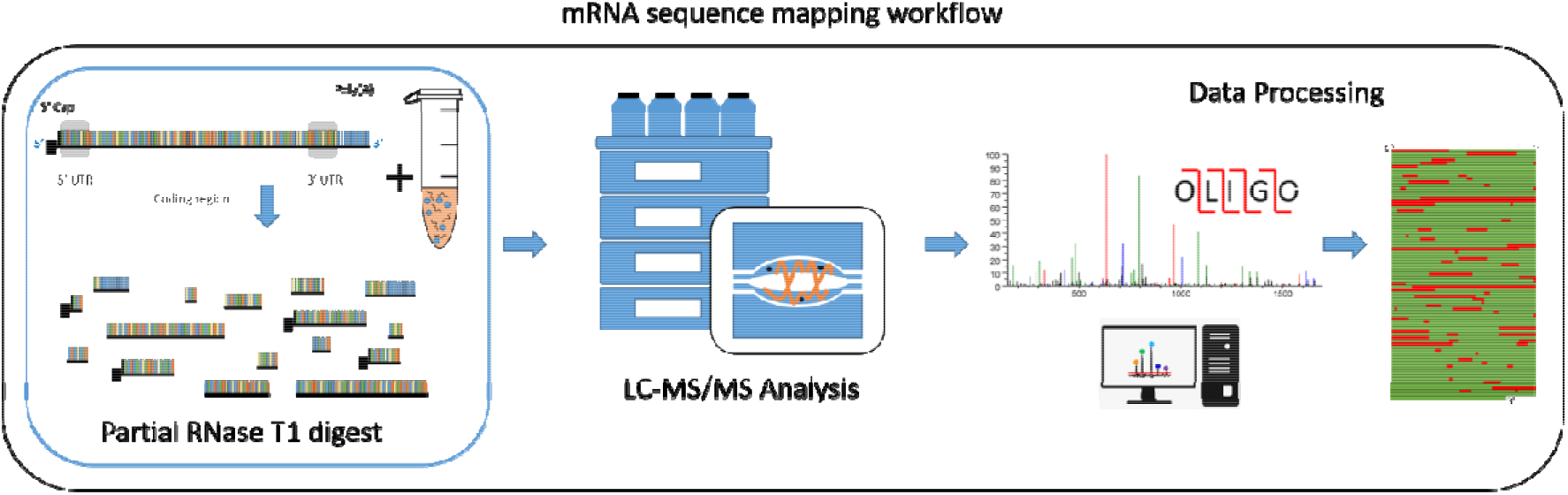
Schematic illustration of mRNA sequence mapping workflow. Partial RNase T1 digests are performed in conjunction with LC MS/MS analysis for automated oligoribonucleotide annotation and identification prior to sequence mapping.

Manual analysis of digested RNA and mRNA LC MS/MS data is often a complex and time-consuming process, highlighting the need for automated software tools to support data interrogation and oligoribonucleotide identification. A number of data analysis tools including SOS ^25^, Ariadne ^26^ and RNAModMapper ^27^ have previously been developed for automated data analysis and identification of oligonucleotide fragmentation in conjunction with the ability for database-matching. Such methods have been utilised in a number of applications for the analysis of complete RNase digests and identification of rRNA modifications. More recently, NucleicAcidSearchEngine (NASE) has been developed to analyse complex oligonucleotide samples containing multiple different RNA modifications in conjunction with statistical validation using false discovery rate (FDR) estimation ^28^.

However, to date, no commercially supported software has been developed that can readily analyse the complex LC MS/MS data generated from partial RNase digests, which contain large numbers of multiple missed cleavages generated from large mRNA molecules. Here we describe the implementation of BioPharma Finder software for oligoribonucleotide identification and the subsequent sequence mapping of partially digested mRNA samples. The data analysis software provides automated tools for the identification of chromatographic components within an RNA or mRNA sample digest. The monoisotopic mass as well as the MS/MS fragmentation pattern of the identified components are compared to the predicted oligoribonucleotide components of the experimental digest. The predicted oligoribonucleotide components are based on a theoretical digest, which includes the prediction of potential missed cleavages produced during the partial digest protocol. Moreover, the software enables the automatic identification of multiple missed cleavages without the requirement for specifying a number of missed cleavages in the search parameters. The systematic approach described here for the identification oligoribonucleotides has been implemented based upon methodologies previously reported for the analysis of therapeutic proteins^29^.

Oligoribonucleotide identification using the data analysis software is based on the evaluation of mass accuracy, isotopic distribution, and charge state determination, as well as through the comparison of experimental and predicted MS/MS fragmentation spectra. The evaluation of mass accuracy, isotopic distribution and charge state determination during component detection allows for the calculation of the monoisotopic mass for each component found within the chromatogram. This enables the examination of each identified oligoribonucleotide across multiple charge states.

The accurate prediction of oligoribonucleotide MS/MS fragmentation is critical in confidently identifying the oligoribonucleotide fragments produced by the mRNA digest. The observed fragmentation spectrum is automatically compared to a predicted fragmentation model generated for each identified sequence. The comparison between experimental and predicted MS/MS fragmentation spectra is utilized to generate a confidence score value based on probability and a similarity match. A high confidence score indicates a good similarity match and that the probability of obtaining a fragment pattern matching the predicted sequence would be low when compared to a random sequence.

In addition, the software automatically calculates an average structural resolution (ASR) value, which provides an indication of the level of fragmentation for each identified oligoribonucleotide. In the ideal case all bonds between each individual nucleotide residue will be broken and resulting fragment ions matched to the predicted MS/MS spectra. A score of 1.0 indicates that each nucleotide bond in the sequence has been fragmented and matched to the predicted MS/MS spectra of the oligoribonucleotide sequence. The combination a high confidence score with low delta mass ppm deviation, coupled with a low ASR value gives a strong confidence of the sequence being correctly matched. Furthermore, the data analysis software also reports the % RNA sequence coverage based on the unique oligoribonucleotides identified and provides powerful visualization tools that show the identified unique oligoribonucleotides mapped to the RNA sequence.

### 3.2. Optimisation of partial RNase T1 digests in conjunction with LC MS/MS analysis

Utilising this new workflow, we performed direct RNA sequence mapping on a number of different mRNA and long RNA, including mRNA corresponding to a modified version of the SARS CoV-2 Spike protein (≈3900 nt), eGFP mRNA (1113 nt), Fluc mRNA (1929 nt) and a control RNA sequence MS2 RNA (3569 nt) (see Supplementary Figure S1). Optimisation of the partial RNase T1 digest was performed by altering the amount of the immobilised RNase T1 and/or the time of the reaction. Figure 2 shows an example of the partial RNase T1 digest performed using varying amounts of the immobilised RNase T1 beads, whilst the amount of RNA, temperature of reaction and reaction time were constant. The results show that as expected increasing the amounts of immobilised enzyme resulted in an increase in the relative abundance of the smaller oligoribonucleotide fragments, eluting earlier and a corresponding decreases in the abundance of the larger oligoribonucleotide fragments, which elute later in the HPLC gradient. In contrast, the lowest amount of immobilised RNase T1 results in an increase in the relative abundance of the larger oligoribonucleotide fragments. Therefore, by simply altering the amount of immobilised RNase T1 enables simple control of the partial RNase T1 digest during optimisation of the direct RNA sequencing workflow. Furthermore, analysis of three replicate partial RNase T1 digests where all digest conditions were constant is shown in Figure 3A. The results show that under these conditions, similar total ion chromatograms (TICs) were generated across the three replicates. To further examine the reproducibility of the partial RNase T1 digests, further analysis of the sequence coverage and unique oligoribonucleotide identifications was assessed. The results show that across the replicate RNase T1 digests, the LC MS/MS analysis the mean unique oligoribonucleotide identifications was 156 (RSD 7.0%) and the mean sequence coverage for the eGFP replicates was 70.6% (RSD 1.7%) under the conditions used (see Figure 3A). These results highlight the reproducible oligoribonucleotide identifications and resulting sequence coverage across the three different replicate partial RNase T1 digests. Further analysis of the retention time stability across the replicates was performed by analysis of selected identified oligoribonucleotides (see Figure 3B). The results show the RSD of the retention time was below 0.3% for each of the oligoribonucleotides shown. These results indicate that the missed cleavages are not generated randomly. RNase T1 will preferentially cleave at accessible single stranded regions within the RNA and as such generate fragments from single stranded loop structures first. Cleavage at sites originally protected in the regions of secondary structure will occur later as the RNA unfolds during digestion. In this way the pattern of oligoribonucleotide fragments produced is reproducible and dependent on the sequence and the secondary structure of the large RNA.

**Figure 2.**
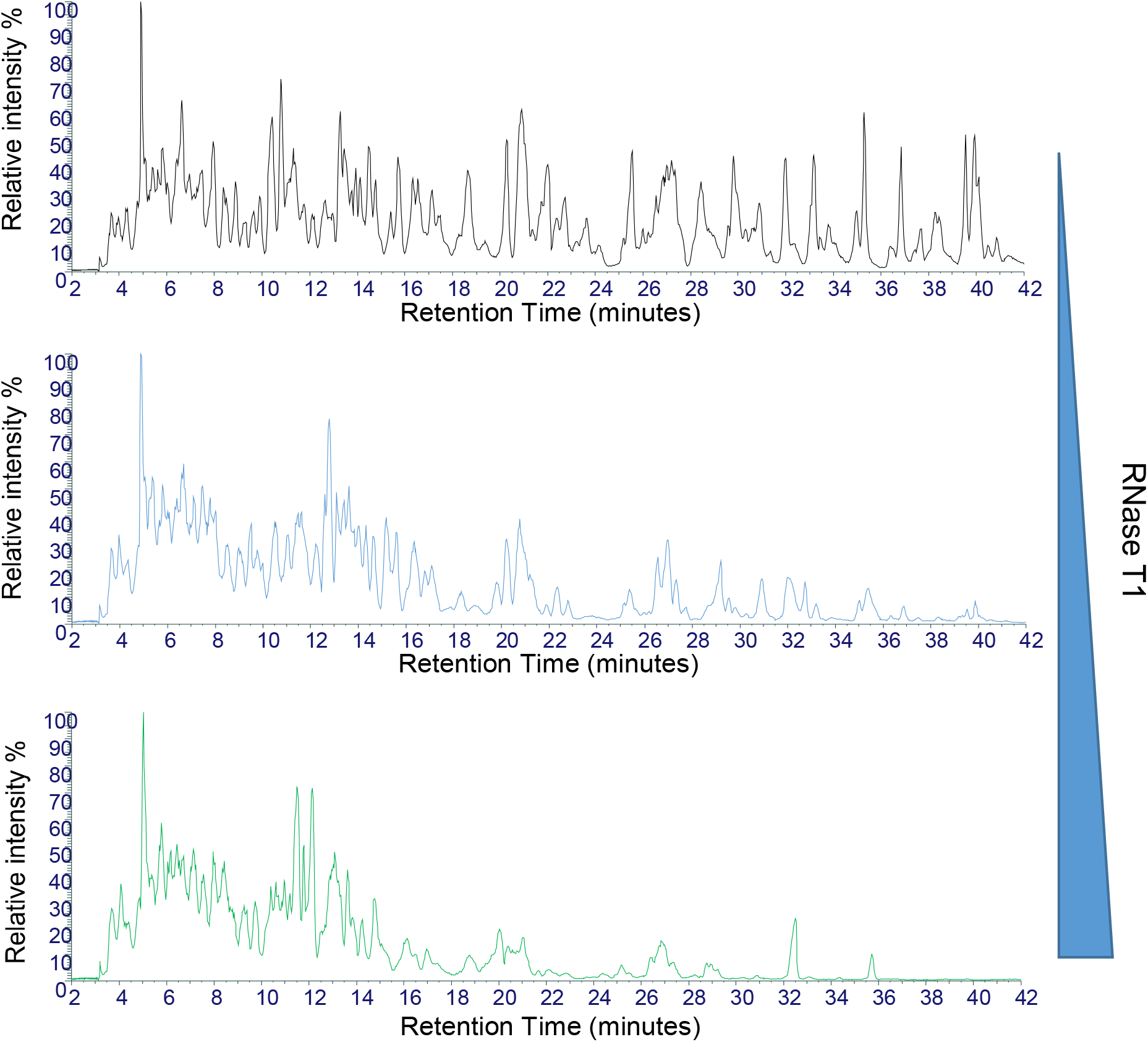
Optimisation of partial RNase T1 digests of mRNA. Total Ion Chromatograms of the partial T1 digests of Fluc mRNA. 20 µg of RNA was incubated with varying amounts of immobilised RNase T1 equivalent to A) 1.25 µl RNase T1 B) 2.5 µl RNase T1 and C) 5 µl RNase T1. All reactions were incubated for 10 mins at 37 °C prior to LC MS/MS analysis.

**Figure 3.**
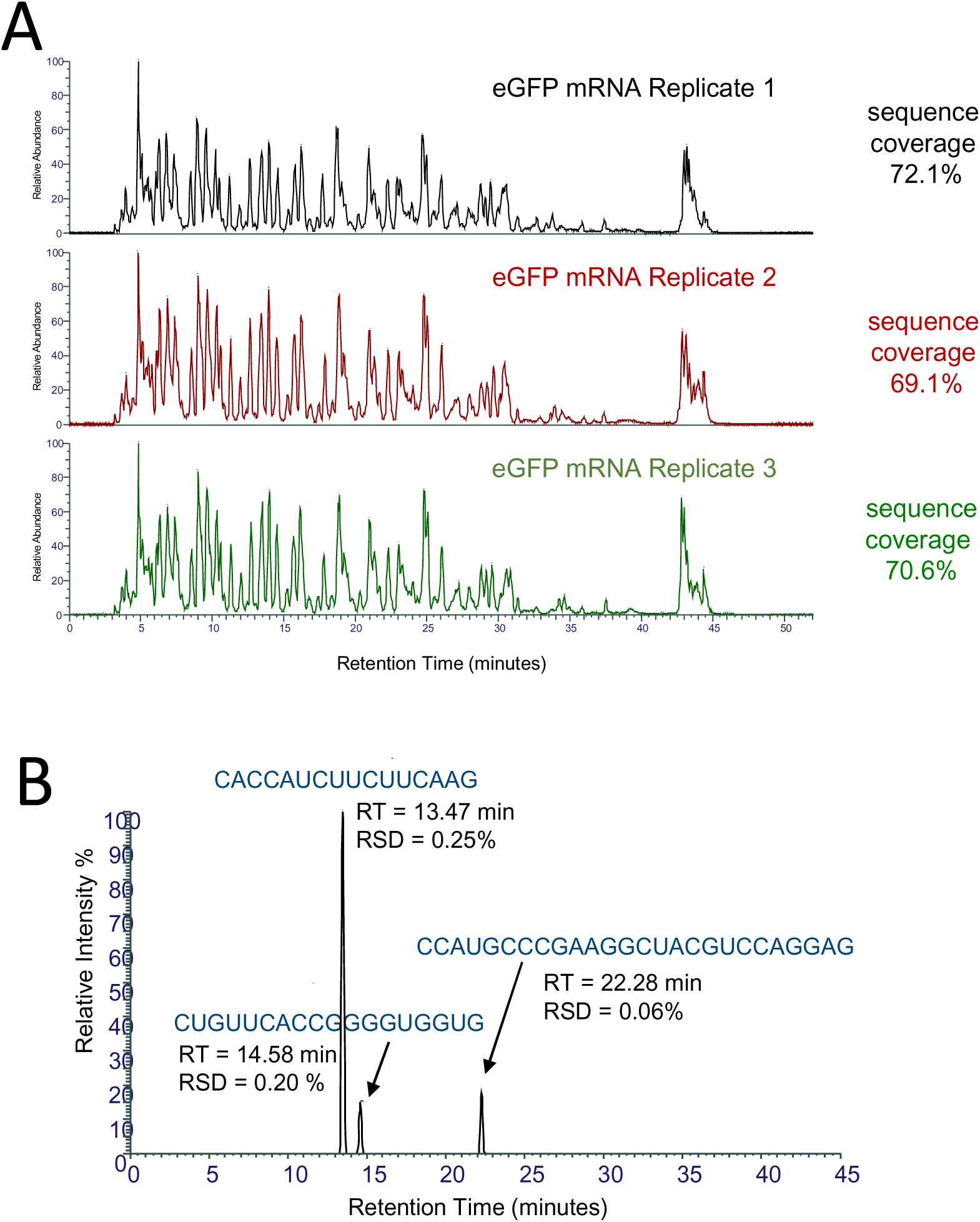
Reproducibility of partial RNase T1 digests of mRNA. A) Total Ion Chromatograms of the partial RNase T1 digests of eGFP mRNA. 20 µg of mRNA was incubated with 2.5 µl of immobilised RNase T1 for 10 mins at 37 °C prior to LC MS/MS analysis. The number of unique oligoribonucleotides and % sequence coverage are highlighted in each replicate. B) Extracted ion chromatogram of three identified unique oligoribonucleotides from the LC MS/MS analysis. The retention time and RSD across the replicates are shown for each oligoribonucleotide.

An example base peak chromatogram (BPC) with selected oligoribonucleotide identifications is shown in Supplementary Figure S2. Using the partial RNase T1 digests, the majority of oligoribonucleotides generated contain a 2’3’-cyclic phosphate termini, with lower amounts of 3’-phosphate termini generated (see Supplementary Table T1/T2). The oligonucleotide fragments elute primarily in size order, with smaller oligoribonucleotides eluting prior to the larger oligoribonucleotides which contained increasing numbers of missed cleavages. The high resolution chromatography is maintained over the useful range of oligonucleotide length from 5 nt to 60 nt and also provides some chromatographic separation of isomeric oligonucleotides. This aids in the accurate identification of the digestion fragments by enabling the collection of diagnostic MS/MS fragmentation ions without interference from overlapping isomers. The gradient is continued to allow the elution of larger fragments and any possible remaining full length mRNA which may be present during preliminary optimisation of the digestion

### 3.3. MS/MS analysis enables identification of sequence isomers and sequence validations across multiple charge states

Mass spectrometry analysis of the partial RNase T1 digests showed that multiple charge states for each oligoribonucleotide were typically observed (see Figure 4). Using the LC MS/MS workflow on the Orbitrap Exploris we were able to generate fragmentation spectra for multiple oligoribonucleotide charge states which increased the confidence of the oligoribonucleotide identifications, enabling sequence validation across multiple charge states (see Figure 4). The use of partial RNase T1 digests also limited the number of smaller oligoribonucleotides produced from the large mRNAs and therefore reduced the number of potential isobaric sequence isomers compared to complete RNase T1 digests. However, a small number were still generated in the analysis of large RNA and mRNA therapeutics, where identification based on accurate mass alone would not enable discrimination. Isobaric oligonucleotides containing the same base composition were typically separated during the chromatography and the MS/MS fragmentation data with the automated sequence annotation enabled identification of the sequence isomers. Examples are shown in Supplementary Figure S3 from the partial RNase T1 digest of the SARS CoV-2 spike protein mRNA.

**Figure 4.**
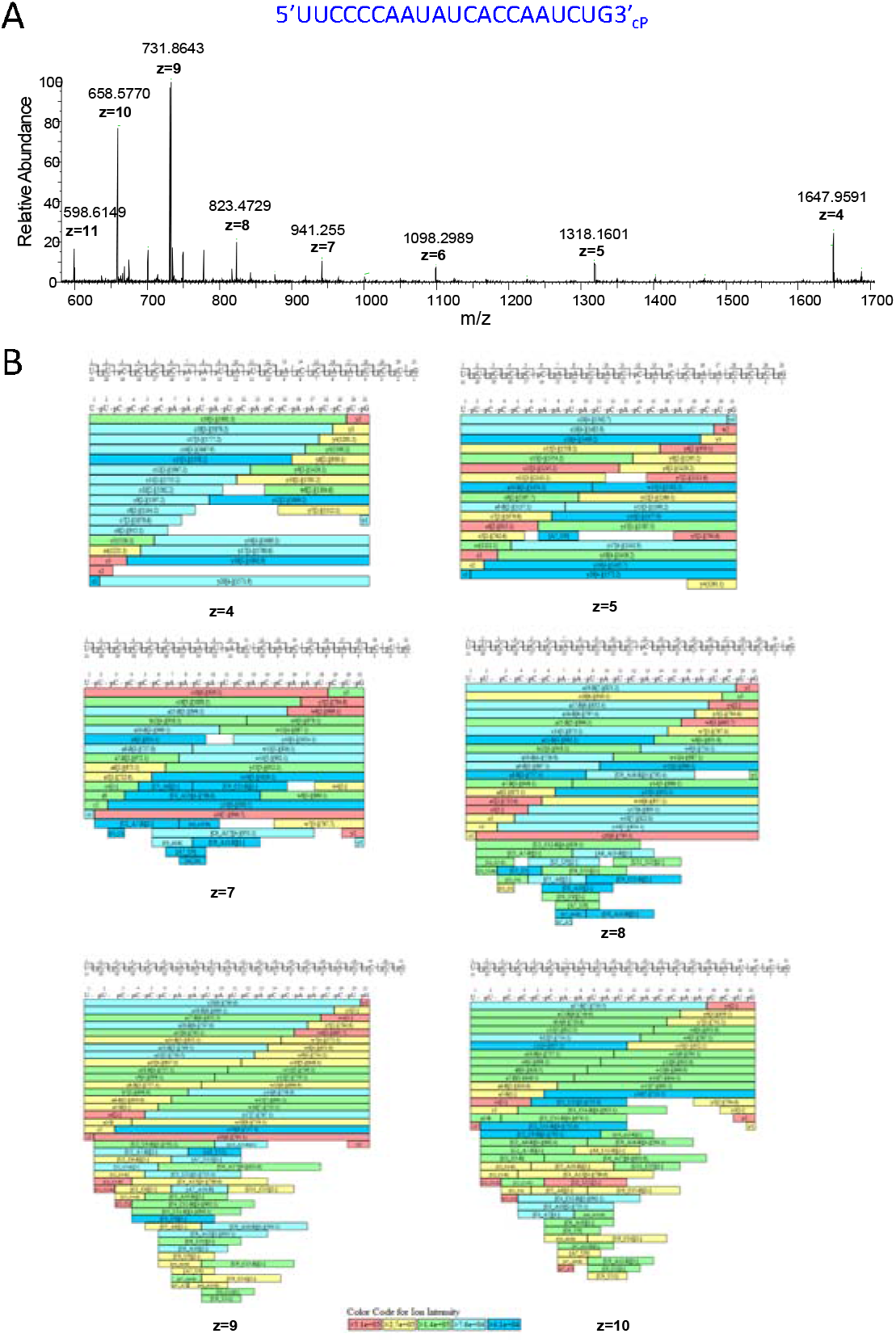
Mass spectrometry analysis of an oligoribonucleotide generated from the partial RNase T1 digest. A) MS spectra of an oligoribonucleotide (5’UUCCC CAAUAUCACCAAUCUG3’-_cP_) from SARS CoV-2 spike protein mRNA. The corresponding monoisotopic m/z and charge states are highlighted. B) Identified oligoribonucleotide fragment ions from the MS/MS spectra are shown for each charge state observed in the MS spectra

### 3.4. Identification of mRNA therapeutics using RNA sequence mapping

Following optimisation of the partial RNase T1 digests, RNA sequence mapping was performed for SARS CoV-2 spike protein mRNA, eGFP mRNA and MS2 RNA using both complete RNase T1 digestions and partial RNase T1 digestions as previously described (see Figure 5A, Supplementary Figure S4). Following LC MS/MS analysis, database searching was performed against the correct RNA sequence and a random RNA sequence of the same GC content and size for each corresponding RNA. The % sequence coverages are shown in Figure 5B and the corresponding sequence coverage maps are shown in Figure 5C. The results show that typically 10-25% sequence coverage based on unique oligoribonucleotide identifications (identified sequences located to a specific unique position on the RNA) was obtained with a complete RNase T1 digestion, consistent with previous RNase sequence mapping of large RNA ^14^. Moreover, as expected, large numbers of non-unique oligoribonucleotides were identified using complete RNase T1 digestion of the large RNA. In contrast, analysis of partial RNase T1 digests revealed significantly higher sequence coverage of SARS CoV-2 spike protein and eGFP mRNA, 86.8 and 86.2% respectively (see Figure 5B). Sequence coverage of the eGFP mRNA not including the polyA tail was 95.9%.

**Figure 5.**
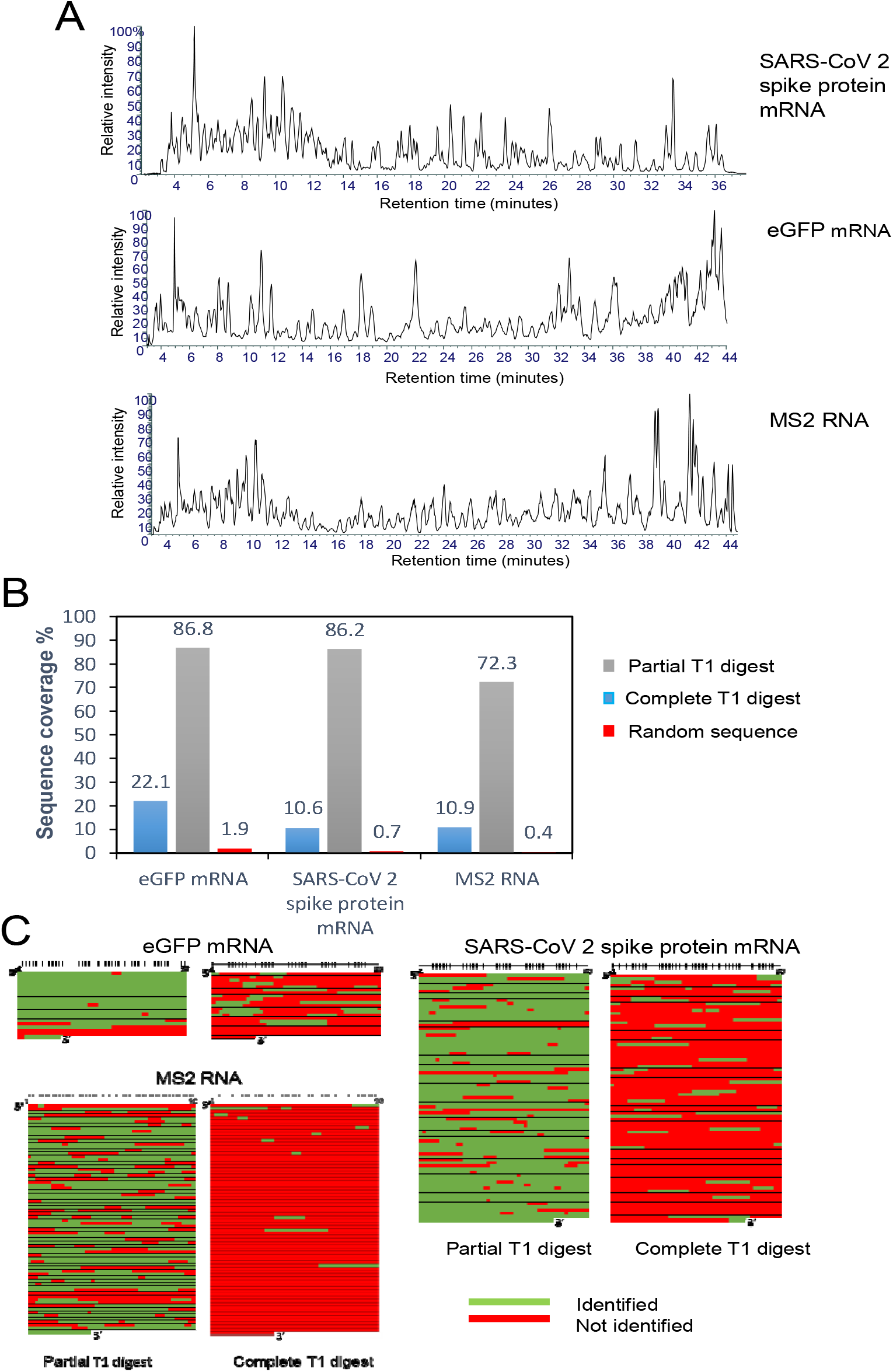
RNA sequence mapping of mRNA therapeutics and long RNA. A) Total Ion Chromatograms of the partial RNase T1 digests of RNA. In the RNase T1 digests, 20 µg of RNA was incubated with 2.5 µl of immobilised RNase T1 for 10 mins at 37 °C. B) Bar chart shows the % sequence coverage obtained for the complete RNase T1 digest, partial RNase T1 digest and the partial RNase T1 digest searched against a random sequence of the same size and GC content as the target RNA.

The results show that the partial RNase T1 digests in conjunction with LC MS/MS analysis is able to identify with high sequence coverage each of the corresponding mRNA from a single analysis. Based on the identification of higher number of larger unique oligoribonucleotides, a significant increase in sequence coverage of the mRNA is obtained using partial RNase T1 digests in comparison to the complete RNase T1 digest. Importantly, for each data set analysed there was very low or no matches against a random RNA sequence of the same GC content and size for each corresponding RNA, demonstrating the specificity of this LC MS/MS method in conjunction with the parameters used in the oligoribonucleotide identifications and sequence mapping software.

Further RNase sequence mapping of a large RNA was performed using the 3569 nt RNA from MS2 phage. This RNA molecule was used as a standard RNA and is a challenging RNA molecule to generate sequence information from using RNase mapping, due to the high degree of secondary structure present that is likely to prevent RNase T1 cleaving at double stranded regions within the RNA fragment. Moreover, in the case of partial RNase digests where enzymatic digestion is limited, stable RNAs with high secondary structure elements may be more resistant to RNase cleavage under partial RNase digest conditions. Optimising the partial RNase T1 digest conditions we were able to obtain 72% sequence coverage from a single analysis of this large RNA (see Figure 5A). Furthermore, by combining multiple analyses and varying digestion conditions, it is possible to further increase sequence coverage of the RNA. These results demonstrate that high sequence coverage can be obtained using a simple partial RNase T1 digest on both mRNA therapeutics and long RNA even with high degree of secondary structure present from a single analysis.

### 3.5. Sequence mapping of chemically modified mRNA using LC MS/MS

Two highly efficacious vaccines based on mRNA sequences encoding for a modified version of the SARS-CoV-2 spike protein have recently been developed ^1-2^. The vaccines made by Moderna and Pfizer–BioNTech use mRNA that has been chemically modified to replace the uridine (U) nucleotide with pseudouridine (Ψ). This change is thought to prevent the immune system reacting to the introduced mRNA. To optimize the mRNA structure and reduce its immunogenicity, modified nucleotides including 5-methylcytidine, pseudouridine, N1-methylpseudouridine, 5-methoxyuridine, 5-methyluridine, or N6-methyladenosine have been used. Therefore, in addition to demonstrating the successful direct RNA sequencing analyses of unmodified mRNA using partial RNase T1 digests in conjunction with LC MS/MS analysis, further work was performed using chemically modified mRNA. mRNA corresponding to the Fluc sequence containing either uridine or 5-methoxyuridine (replacing all uridines) was analysed using the workflow previously described and the resulting TICs are shown in Figure 6A. For data analysis, 5-methoxyuridine was added using the sequence editor in BioPharma Finder to generate a new sequence where all uridines were replaced by 5-methoxyuridine which was subsequently used in the data analysis. The results show that the LC MS/MS analysis resulted >90% sequence coverage of the unmodified Fluc mRNA (ORF) from a single analysis, consistent with previous data (see Figure 6B). Moreover, using the same workflow >90% sequence coverage of the modified Fluc mRNA (5-methoxyuridine) was obtained demonstrating the ability to generate high sequence coverage of chemically modified mRNA and detect chemically modified oligoribonucleotides in the LC MS/MS analysis. Further analysis and manual validation of the MS/MS spectra was also performed. Figure 6C shows the MS/MS spectra of an oligoribonucleotide generated from the partial RNase T1 digest from both the unmodified mRNA and the same corresponding oligoribonucleotide from the 5-methoxyuridine mRNA. The specific fragment ions corresponding to sites of the 5-methoxyuridine modifications are highlighted in red. Further control analysis was performed by searching the LC MS/MS data from the chemically modified mRNA partial digest against the unmodified Fluc mRNA sequence, in this case as expected only oligoribonucleotides which do not contain uridine were identified.

**Figure 6.**
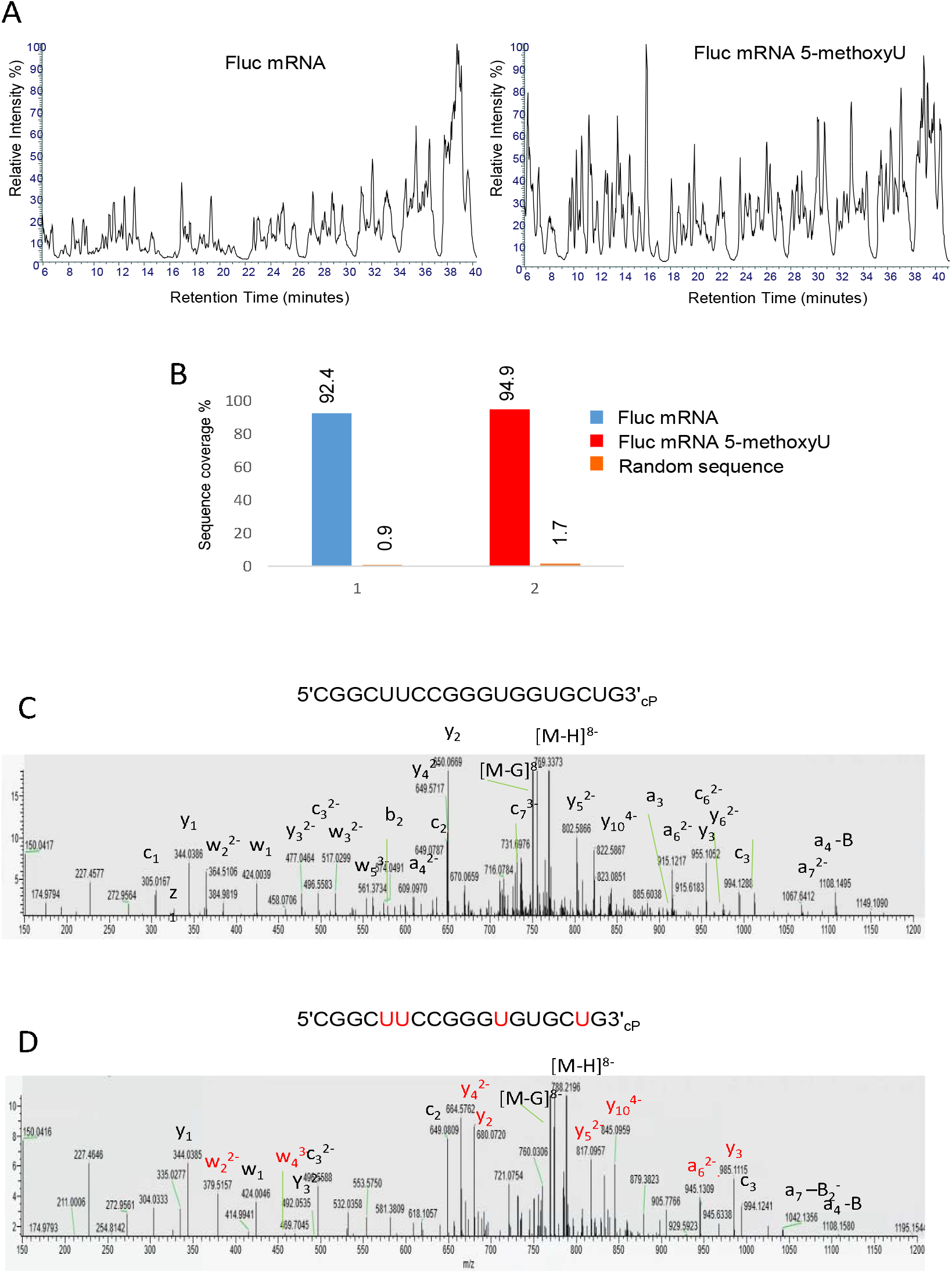
RNA sequence mapping of chemically modified mRNA. A) Total Ion Chromatograms of the partial RNase T1 digests of Fluc mRNA and Fluc 5-methoxyU mRNA. 20 µg of RNA was incubated with 1.25 µl of immobilised RNase T1 for 10 mins at 37 °C prior to LC MS/MS analysis. B) Bar chart showing the % sequence coverage of the partial RNase T1 digest of Fluc mRNA (ORF sequence), chemically modified mRNA (ORF) and random RNA sequence. D) MS/MS spectra of the oligoribonucleotide CGGCUUCCGGGUGGUGCUG_cP_ and the corresponding oligoribonucleotide where the uridines are replaced with 5-methoxyuridines. The corresponding fragment ions are highlighted and those fragment ions specific for the 5-methoxyuridine oligoribonucleotide are highlighted in red.

### 3.6. Identification of RNA impurities or mixed RNA samples using RNA sequence mapping

Further experiments were performed using the partial RNase T1 digests in conjunction with LC MS/MS analysis in an approach to identify the presence of potential impurities in mRNA samples and the ability to detect mRNA in mixed RNA samples. SARS-CoV-2 spike protein mRNA was mixed at varying mass ratios with rRNA which was used to represent potential impurities prior to partial RNase T1 digests and LC MS/MS analysis (see Supplementary Figure S5). The results show that using ratios of 10:1 (mRNA:rRNA) we were able to detect the presence of the rRNA in the mRNA sample. These results demonstrate the ability of LC MS/MS method to detect low-level impurities of a known RNA sequence within an mRNA sample.

## 4. CONCLUSIONS

Partial RNase digestions using RNase T1 immobilised on magnetic particles in conjunction with high resolution tandem mass spectrometry analysis with automated oligoribonucleotide identifications enabled >80% sequence of coverage of a range of large RNAs and mRNA therapeutics from a single analysis. This novel approach demonstrated significant improvements in sequence coverage compared to conventional complete RNase T1 digestions. The automated data analysis enabled rapid verification of the long RNA sequences from complex oligoribonuceotide LC MS/MS data sets. Furthermore, high sequence coverage with no or low sequence matches against random control RNA sequences was obtained, demonstrating the specificity of the analytical workflow in conjucntion with the parameters used for RNA sequence mapping. mRNA therapeutics have emerged as a new important class of therapeutic and require the development of new analytical methods to analyse mRNA critical quality attributes and confirm identity. Direct sequencing of the mRNA is now possible in a simple automated workflow using this new approach. The analytical workflow, including automated sample preparation can be completed within 90 minutes. Simple partial RNase T1 digestion with LC MS/MS analysis and automated data analysis offers a rapid, high throughput method for the analysis of mRNA therapeutics and the ability to analyse important mRNA critical quality attributes including RNA sequence integrity and RNA sequence identity. Furthermore, the same workflow can be used for sequence mapping of chemically modified mRNA and to identify impurities present in the mRNA. Further development of simple assays for quality control testing of mRNA vaccines could be developed utilising similar workflows.

## ASSOCIATED CONTENT

IP-RP HPLC analysis of intact SARS CoV-2 spike protein mRNA, eGFP mRNA and MS2 RNA (Figure S1). Base peak chromatograms of partial RNase T1 digests of eGFP and SARS CoV-2 spike protein mRNA with identified oligoribonucleotide fragments (Figure S2). Identification of sequence isomers from the partial RNase T1 digest of the SARS CoV-2 spike protein mRNA (Figure S3). Total ion chromatograms of the complete RNase T1 digests of SARS CoV-2 spike protein mRNA, eGFP mRNA and MS2 RNA (Figure S4). Analysis of SARS CoV-2 spike protein mRNA samples with rRNA impurities at 1:10 and 1:100 mass ratios (Figure S5). All oligonucleotide identifications from BioPharma Finder are shown in Supplementary Tables T1/T2.

## ACKNOWLEDGEMENTS

This work was supported by funding from the Engineering Physical Science Research Council EPSRC Impact Acceleration Account (EP/R511754/1) to the University of Sheffield. M.J.D. acknowledges further support from the Biotechnology and Biological Science Research Council (BB/M012166/1).

