## Supplementary Figures for "Characterisation and sequence mapping of large RNA and mRNA therapeutics using mass spectrometry"

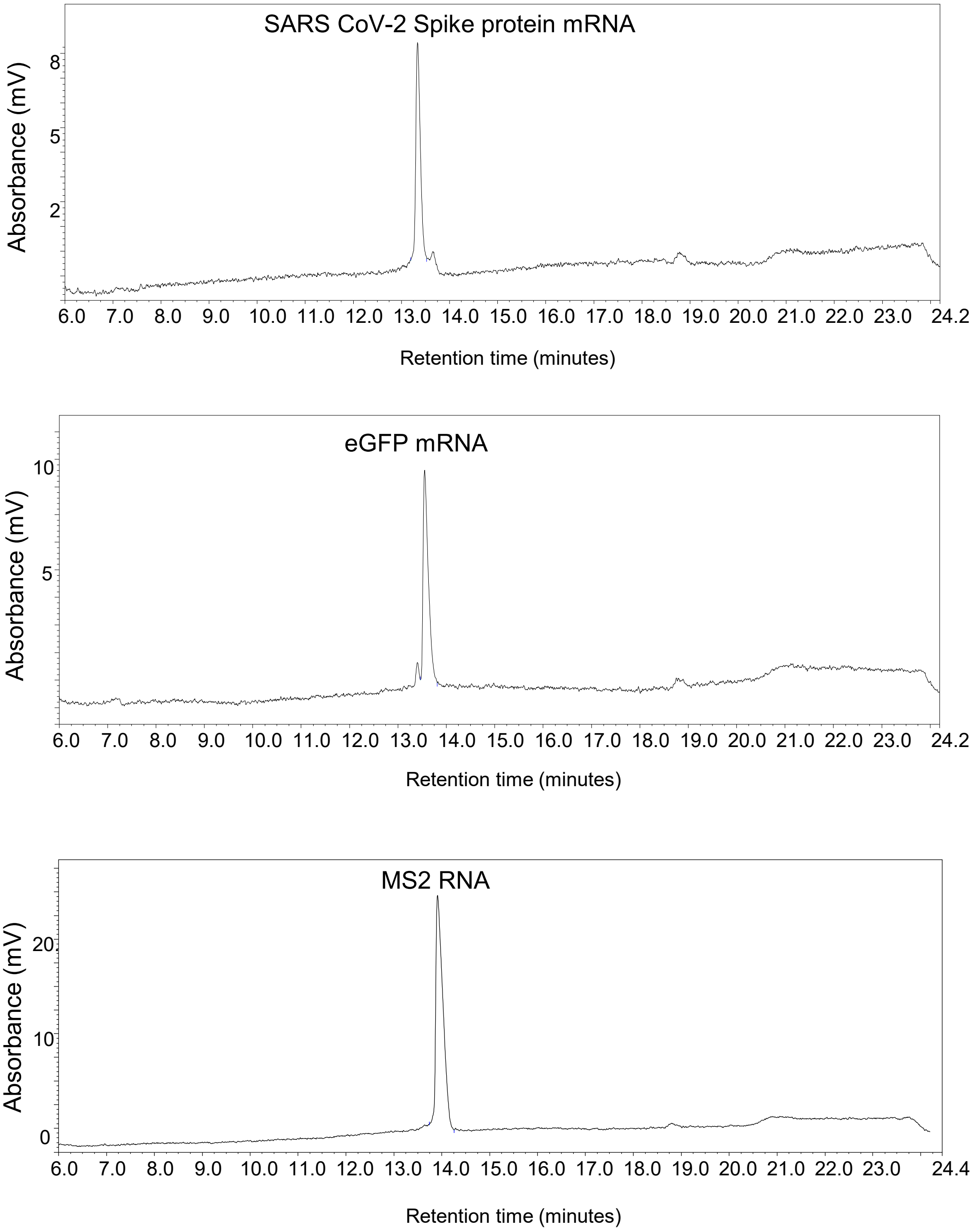


**Supplementary Figure S1**. IP RP HPLC analysis of intact RNA. A) Chromatogram shows the analysis of the SARS CoV-2 Spike protein mRNA following synthesis using IVT and purification. B) Chromatogram show the analysis of the eGFP mRNA following synthesis using IVT and purification. C) Chromatogram of the MS2 RNA. 100 ng of each RNA was analysed using IP RP HPLC with UV detection at 260 nm.

Supplementary Figure S1

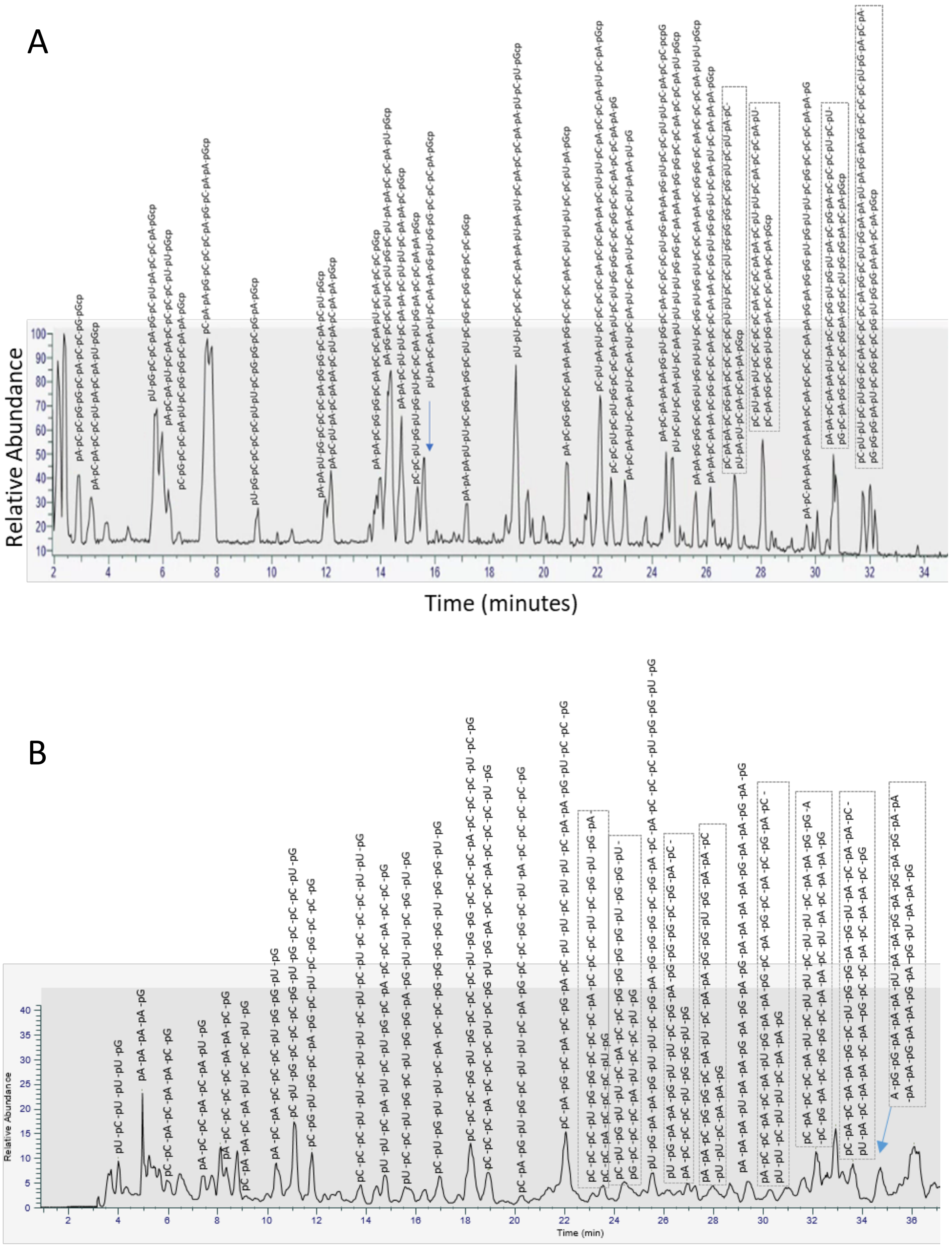


**Supplementary Figure S2**. Base Peak Chromatograms of the partial RNase T1 digests of RNA. IP RP HPLC in conjunction with MS analysis was used to analyse partial RNase T1 digests of A) SARS CoV-2 spike protein mRNA. B) eGFP mRNA. Selected identified oligoribonucleotides are highlighted.

Supplementary Figure S2

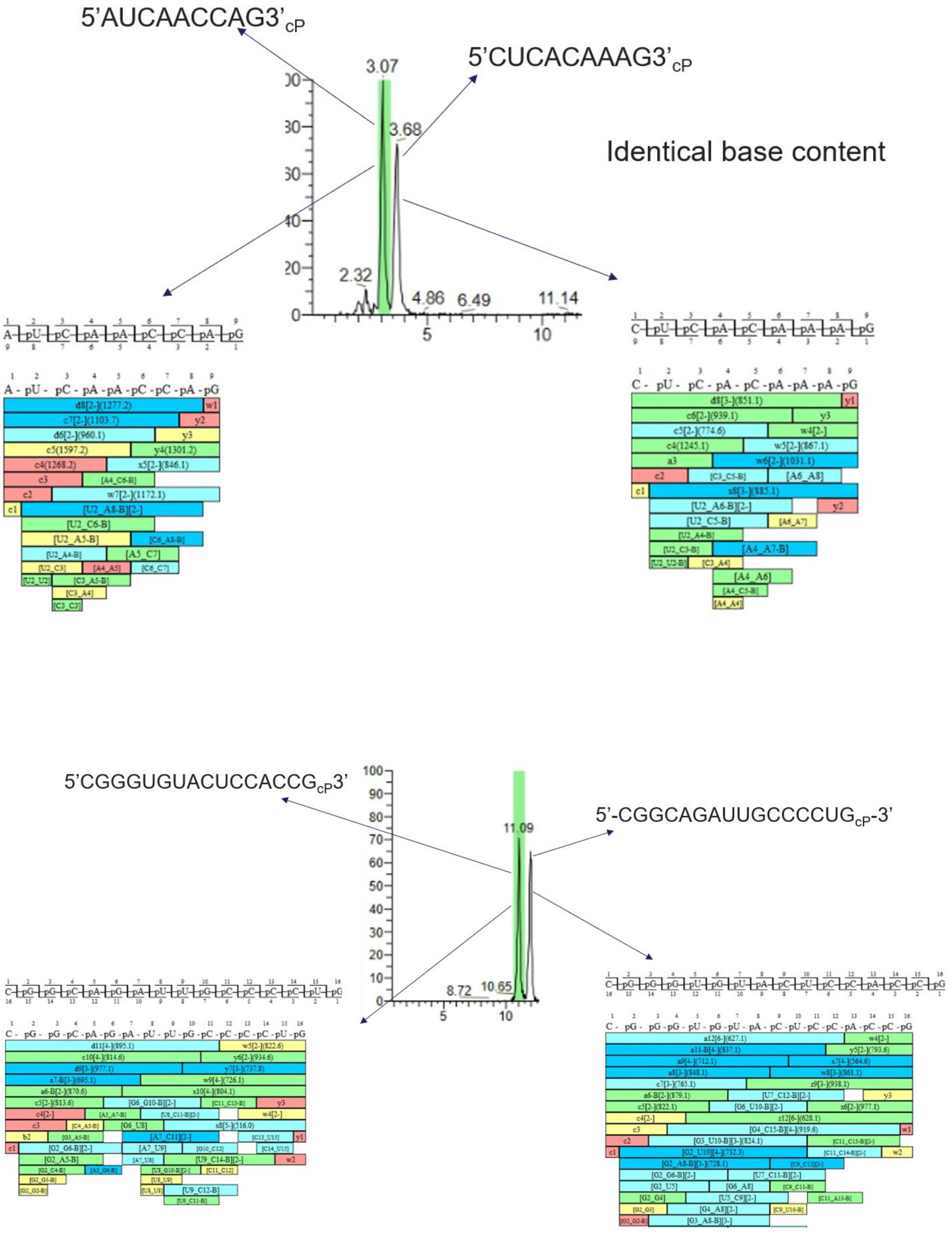


**Supplementary Figure S3**. Identification of sequence isomers from the partial RNase T1 digest of the SARS CoV-2 spike protein mRNA. Isobaric oligonucleotides containing the same base composition were typically separated during the chromatography and the $MS^2$ fragmentation data with the automated sequence annotation enabled identification of sequence isomers.

Supplementary Figure S3

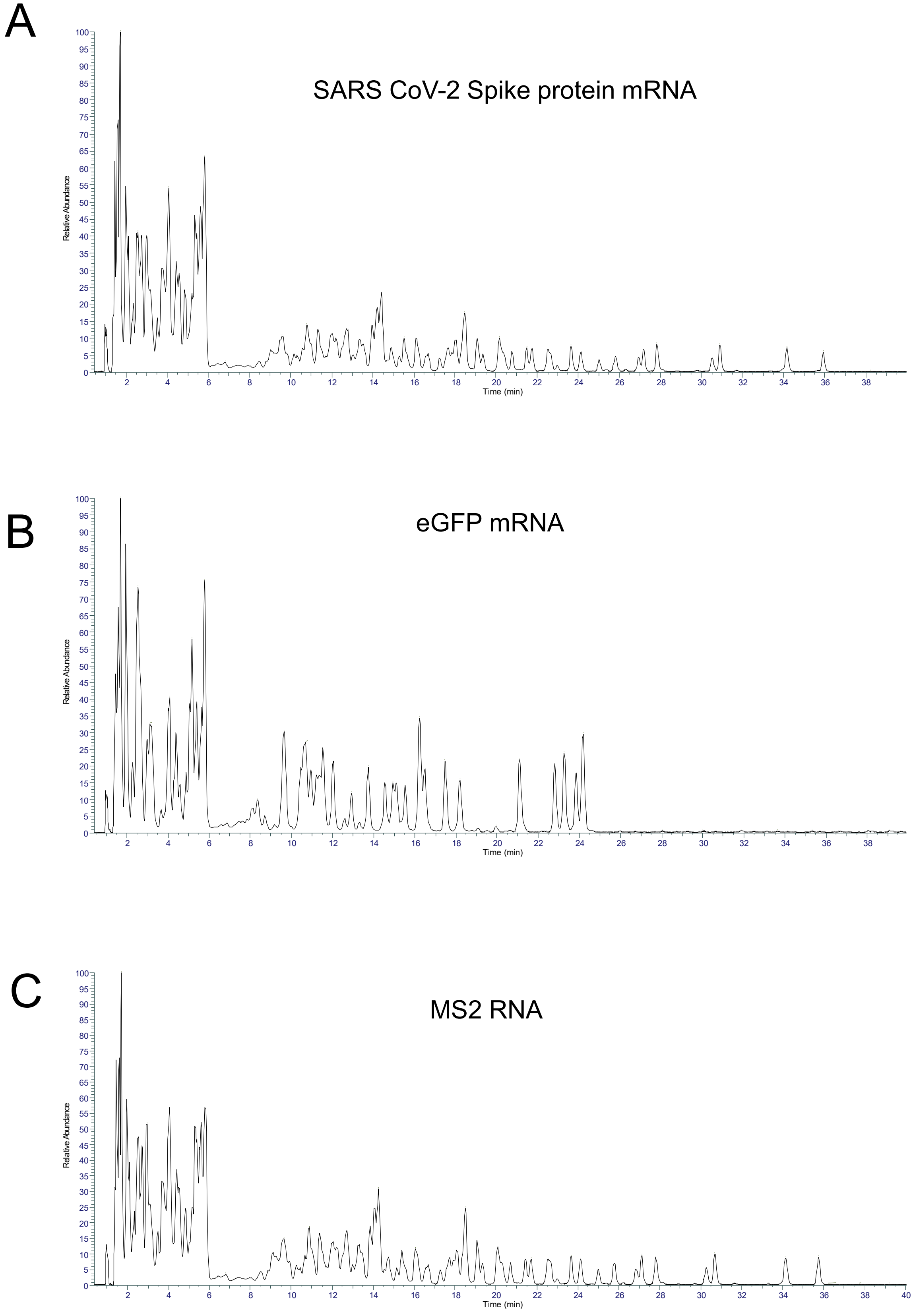


**Supplementary Figure S4**. Total Ion Chromatograms of the complete RNase T1 digests of RNA. IP RP HPLC in conjunction with MS analysis was used to analyse complete T1 digests of A) SARS CoV-2 spike protein mRNA B) eGFP mRNA and C) MS2 RNA. In each RNase T1 digest 10 µg of RNA was incubated with RNase T1 for 4 hours at 37 °C.

Supplementary Figure S4

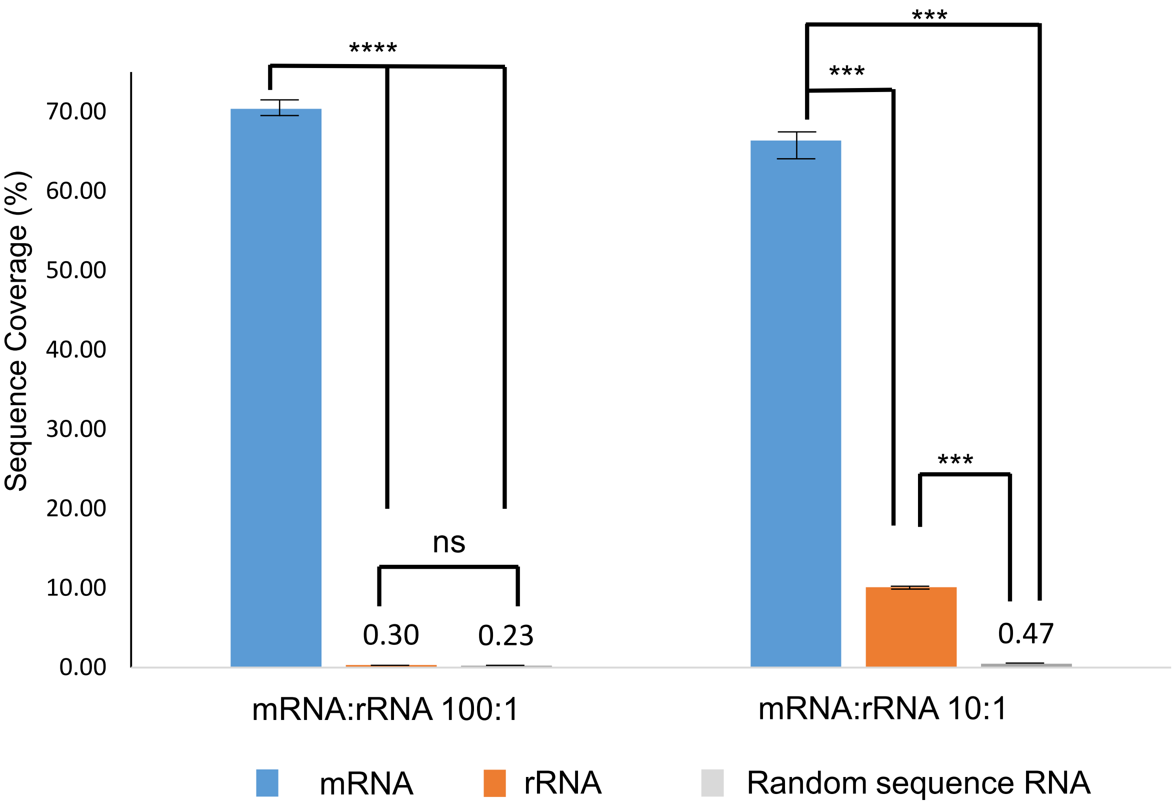


**Supplementary Figure S5**. Bar chart of the % sequence coverage of the SARS CoV-2 spike protein mRNA, rRNA and random sequence RNA. The mRNA and rRNA were mixed in either 100:1 or 10:1 mass ratio prior to partial RNase T1 digestion and LC MS/MS analysis. In all graphs, mean and S.E. are plotted (n=3). Stars denote significance as calculated by T-tests. ns = P > 0.05, * = P ≤ 0.05, ** = P ≤ 0.01, *** = P ≤ 0.001, **** = P ≤ 0.0001

Supplementary Figure S5
